# Sequential involvements of macaque perirhinal cortex and hippocampus in semantic-like memory including spatial component

**DOI:** 10.1101/2022.08.15.504057

**Authors:** Cen Yang, Yuji Naya

## Abstract

The standard consolidation theory suggests the critical involvement of the hippocampus (HPC) in acquiring new knowledge, while the perirhinal cortex (PRC) is involved in its long-term storage (i.e., semantic memory). Converging studies have shown exclusive involvement of the PRC in item processing, while the HPC relates the item with a spatial context. These two lines of literature raise the following question; which brain region is involved in semantic recall that includes the spatial components? To solve this question, we applied an item-location associative (ILA) paradigm in a single-unit study using non-human primates. We trained two macaques to associate four visual item pairs with four locations on a background map before the recording sessions. In each trial, one visual item and the map image at a tilt (−90 to 90 degrees) were sequentially presented as the item-cue and the context-cue, respectively. The macaques chose the item-cue location relative to the context-cue by positioning their gaze. Neurons in both PRC and HPC but not area TE exhibited item-cue responses which signaled retrieval of item-location associative memory. This retrieval signal first appeared in the PRC before appearing in the HPC. We examined whether neural representations of the retrieved locations were related to the external space where the macaques viewed. A positive representation similarity was found in the HPC but not PRC, suggesting a contribution of the HPC to relate the retrieved location with a first-person perspective of the subjects. These results suggest their distinct but complementary contributions to semantic recall including spatial components.

## INTRODUCTION

Semantic memory, the memory of factual knowledge, is a subcategory of declarative memory (Squire and Zola-Morgan, 1991; Tulving, 1972). Semantic memory is acquired through repetitive experiences that share common facts in different spatiotemporal contexts. This common fact would eventually be stored as semantic memory separately from a unique spatiotemporal context in each experience. This contrasts with another subcategory of declarative memory (i.e., episodic memory), which allows us to re-experience a particular past event (Tulving, 1972). Therefore, semantic memory is essentially formed in an allocentric manner, while episodic memory accompanies the unique spatiotemporal contexts to reconstruct a particular personal event in the “self-referenced” manner or the “first-person perspective” (Buzsáki et al., 2022; Buzsaki and Moser, 2013; Naya, 2016b; Schacter et al., 2007; Tulving, 1972). However, when we recall semantic knowledge, particularly that contains spatial components, we may bring it to mind from a first-person perspective. Assume a scenario where someone asks where the Statue of Liberty is in the United States; when a map is presented, even if at a tilt, the Statue of Liberty can be easily located on it (Fig. 1A), which would suggest an allocentric representation of semantic memory. In the absence of the map, you may get mental imagery of the retrieved location to represent it from a first-person perspective (e.g., rightward) by assuming a particular spatial context (e.g., top for the north).

**Fig. 1.**
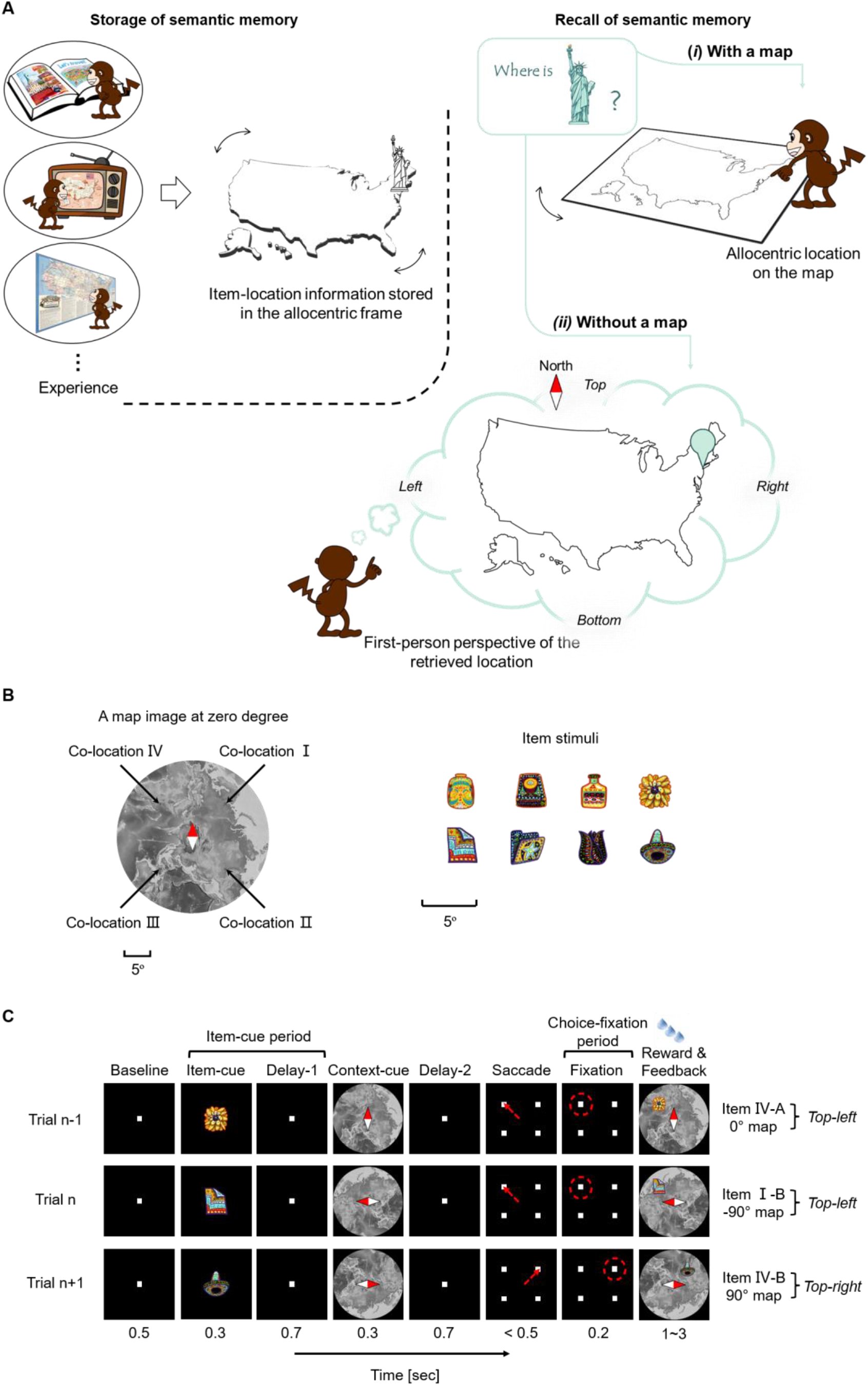
Semantic memory including spatial components. **(A)** (Left) Through repetitive experiences (e.g., reading a book in a study room, etc.), item-location associations (e.g., Statue of Liberty ↔ northeast in the US) can be stored in the allocentric frame. The stored memory, which does not accompany a unique spatiotemporal context from a single episode (e.g., watching TV in a living room), could be categorized into semantic memory. (Right) The recall process of the item-location associative memory may differ between the presence (*i*) and absence (*ii*) of a map on which the item’s location can be shown. (*i*) When the map is present, the item can be located on it no matter how it is tilted (0° to 360°), suggesting the semantic recall in the allocentric frame. (*ii*) In the absence of the map, the item can be located in an assumed spatial context (e.g., top for the north), providing mental imagery of the retrieved location from the first-person perspective. **(B)** Item-location association (ILA) pattern. Two items, one from set A (e.g., I-A) and the other from set B (e.g., I-B), were assigned to each location (e.g., co-location I) on the map image. Scale bars for both item and map stimuli, with a 5° visual angle. **(C)** Schematic diagram of the ILA task. An item-cue and a context-cue were sequentially presented in each trial. The monkeys’ gaze was fixated on the center until the end of the delay-2, then saccade to the target location (indicated by the red arrowhead and red dashed circle) according to the two cues. A successful trial was rewarded with juice paired with feedback showing the associated location of the item-cue on the context-cue. Relative sizes of the stimuli were magnified for display purposes.

Concerning the responsible brain regions for the two subcategories of declarative memory, previous neuropsychological studies have indicated the critical involvement of the hippocampus (HPC) in encoding both new episodic memory and semantic memory, whereas HPC lesions impair recollection of episodic memory but spare retrieval of semantic memory (Milner, 1959; Moscovitch et al., 2006; Nadel and Moscovitch, 1997; Squire and Wixted, 2011). In contrast, the anterior temporal lobe, particularly the perirhinal cortex (PRC), is involved in conceptual processing, which depends on factual knowledge stored as semantic memory. The functional dissociation between the PRC and the HPC is also suggested by a series of dual-process models in an item recognition paradigm (Davachi, 2006; Eichenbaum et al., 2007; Mayes et al., 2007); the PRC contributes to recognition by providing item familiarity, whereas the HPC contributes to recognition by recollecting the spatiotemporal context in which the item was presented. These models are consistent with anatomical evidence suggesting separate information processing along the ventral path-PRC stream and along the dorsal path-parahippocampal cortex (PHC) stream and their relational processing in the HPC (Chen and Naya, 2021; Haxby et al., 1991; Lavenex and Amaral, 2000; Ranganath and Ritchey, 2012). The distinct mnemonic functions of the PRC and the HPC in these two lines of preceding literature (“semantic memory vs. episodic memory” and “familiarity/item vs. recollection/relation”) raise the question of which brain area is involved in the retrieval of semantic memory, particularly when it contains spatial components.

To address this question, we used the item-location association (ILA) paradigm in macaque electrophysiology (Yang and Naya, 2020). In this study, before recording sessions, we first repeatedly trained monkeys to associate visual items with locations on a background map image (Fig. 1B and Fig. S1). Because the map image was presented at a tilt with a random orientation, the monkeys needed to store a common relationship between the items and corresponding locations relative to the map image across the past trials rather than to store a unique spatiotemporal context in each trial. Thus, the ILA paradigm could potentially be a useful animal model to test semantic-like memory. In each trial, one of the learned visual items was presented as an item-cue, and then the randomly tilted map image was presented as a context-cue. The monkeys answered the location associated with the item-cue relative to the context-cue by positioning their gaze (Fig. 1C). We recorded spike firings of single neurons during the monkeys performing the ILA task. We examined neuronal responses to the item-cues in the absence of the map image (i.e., before the context-cue), which might remind the monkeys of locations associated with the item-cues. To evaluate mnemonic components carried by neural responses to the item-cue, we used two types of measures. In the first measure, we tested the presence/absence of item-location association effect by examining neurons’ responses to pairs of the item-cues (“co-locating items”) assigned to the same locations (“co-locations”) on the map image. In the second measure, we evaluated a relationship between neural representations of the retrieved locations during the item-cue period, and those of external space coupled with the gaze positions during the monkeys answered target locations (choice-fixation period). The first measure revealed an earlier appearance of the item-location associative effect in the PRC before HPC. The second measure indicated that neural representations of retrieved item-cue locations were related with those of external space where the monkeys viewed, only in the HPC but not in the PRC. This relationship was observed only when it was assumed that a spatial context given by a particular context-cue would accompany the recollection of the item-cue location. These results suggest that the PRC and the HPC are involved in remembering semantic memory in distinct manners, at least when spatial components are contained. In addition to the PRC and HPC, we recorded signals from area TE (TE) of the ventral pathway and the PHC as control brain regions.

## RESULTS

Two rhesus macaques were trained to perform the ILA task. In the ILA task, four visual item pairs (“co-locating items”) were assigned to four different locations (“co-locations”) on an image of a map (Yang and Naya, 2020). The same eight visual items and one map image were used during all the recording sessions (Fig. 1B). In each trial, a randomly chosen visual item was presented as an item-cue. Then the map image was presented at a tilt with a randomly chosen orientation (−90° to 90°, 0.1° step) as a context-cue (Fig. 1C). By moving their gaze positions, the monkeys reported a target location (e.g., *Top-left* in the trial n, Fig. 1C), which corresponded to the relative location of the co-location of the item-cue (e.g., *co-location I*) on the tilted map image of the context-cue (e.g., *-90°*). While the monkeys performed the ILA task, we recorded single-unit activity from a total of 1,175 neurons from the four brain areas (Fig. 2A, Table S1). During the recording session, the orientation of the context-cue was pseudorandomly chosen from among five orientations (−90°, −45°, 0°, 45°, and 90°). The task was correctly performed by both monkeys (chance level = 25%) at rates of 80.5% ± 8.4% (mean ± standard deviation; Monkey B, n = 435 recording sessions) and 95.1% ± 5.8% (Monkey C, n = 416 recording sessions). We used only correct trials for the following analyses. The HPC data was partly reported in a previous study (Yang and Naya, 2020).

**Fig. 2.**
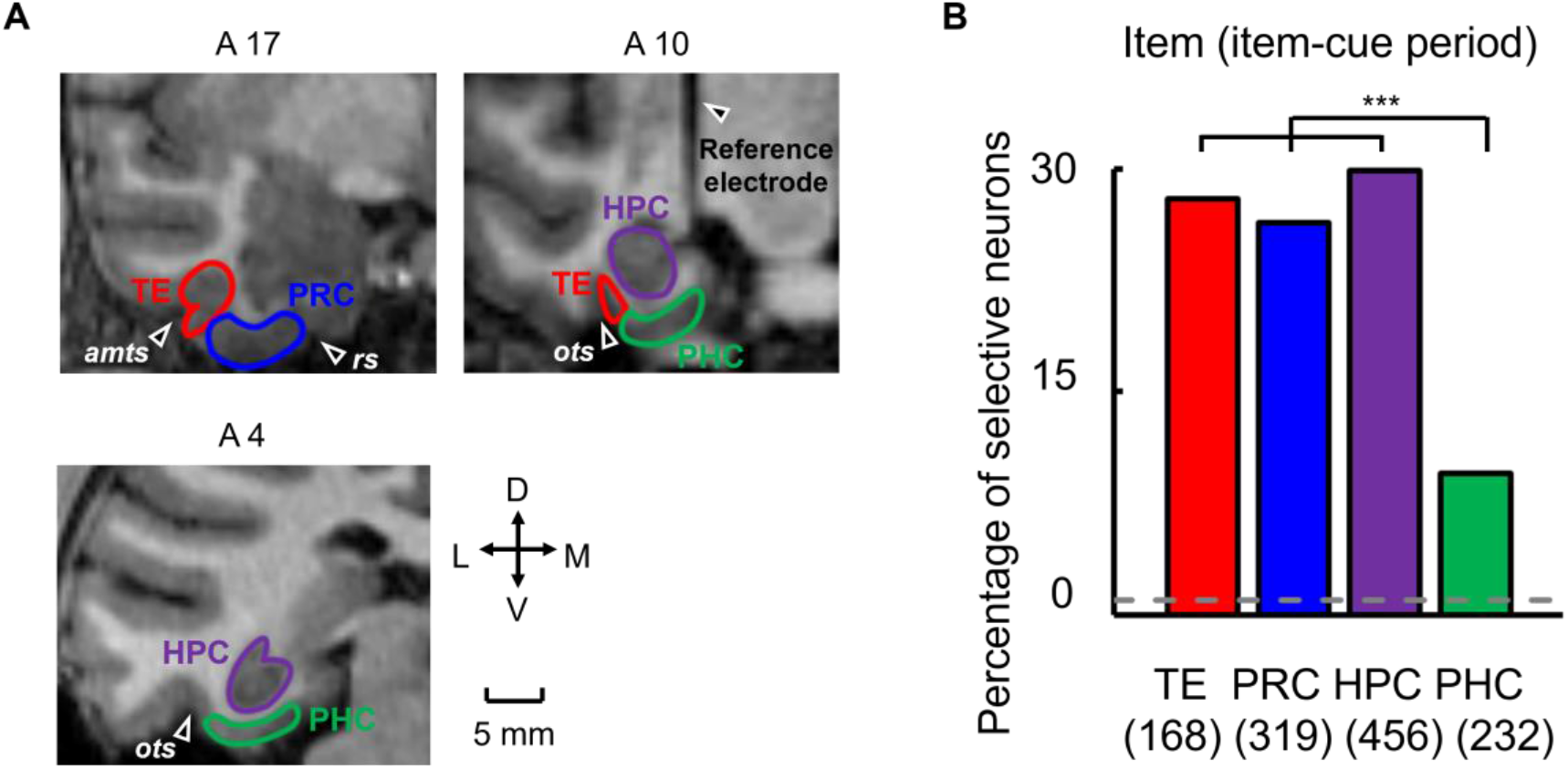
Item-cue selective activities in the MTL and TE. **(A)** Recording regions. MRI images corresponding to the coronal planes anterior 4, 10, and 17 mm from the interaural line of monkey C (right hemisphere). The recording regions are the perirhinal cortex (PRC), hippocampus (HPC), and parahippocampal cortex (PHC) of the MTL and TE. A reference electrode implanted in the center of the chamber was observed as a vertical line of shadow in the coronal plane at A10. amts, anterior middle temporal sulcus. rs, rhinal sulcus. ots, occipital temporal sulcus. D, dorsal. V, ventral. L, lateral. M, medial. **(B)** Percentage of item-selective neurons out of recorded neurons in each area. Parentheses, number of recorded neurons in each area. Dashed line, 1.0% chance level. The number of item-cue selective neurons was significantly larger than the chance level (P < 0.0001 for each area, one-tailed binomial test). Asterisk indicates results of a χ-square test: PHC and TE, P < 0.0001***, χ2 = 23.3, d.f = 1; PHC and PRC, P < 0.0001***, χ2 = 24.5, d.f = 1; PHC and HPC, P < 0.0001***, χ2 = 36.0, d.f = 1.

### Retrieval of item-location associative memory

We analyzed the activities of single neurons during the item-cue period, including the presentation of the item-cue (0.3 s) and following delay-1 (0.7 s) periods, to examine their involvement in item-location associative memory. We first selected neurons that exhibited differential item-cue responses among the eight visual items (item-cue selective) at a threshold of *P* < 0.01 (one-way ANOVA). We found a large proportion of the item-cue selective neurons in the PRC (26.3%), TE (28.0%), as well as in the HPC (29.8%) (Fig. 2B, Table S1). In contrast, the proportion of the item-cue selective neurons (9.5%) was significantly smaller in the PHC than in the other three areas (*P* < 0.001 for each area, χ-square test). This result was consistent with previous studies indicating preferential processing of item information in the ventral path-PRC stream (Chen and Naya, 2020b; Haxby et al., 1991; Lavenex and Amaral, 2000; Ranganath and Ritchey, 2012).

Figure 3A shows an example of item-cue selective neurons from the PRC (*P* = 0.0002, *F* (7,78) = 4.78) (see Figs. S2 and S3 for example neurons from TE, HPC, and PHC). This neuron exhibited the largest responses to two item-cues that were assigned to co-location II as well as the second-largest responses to two item-cues that were assigned to co-location III, indicating that similar response patterns were observed for the item-cues with the same co-locations (“co-locating items”). We evaluated the correlated responses to the four pairs of co-locating items by calculating the Pearson correlation coefficient between responses to item-cues ‘I-A’ – ‘IV-A’ and those to item-cues ‘I-B’ – ‘IV-B’ as the “co-location index.” If a neuron showed a pattern of stimulus selectivity that was independent of the items’ co-locations, the expected value of the neuron’s co-location index was ‘zero.’ The co-location index of the example neuron shown in Fig. 3 was significantly positive (*r* = 0.98, *P* = 0.0024, two-tailed permutation test), suggesting an effect of the item-location associative memory on its response pattern to the item-cues.

**Fig. 3.**
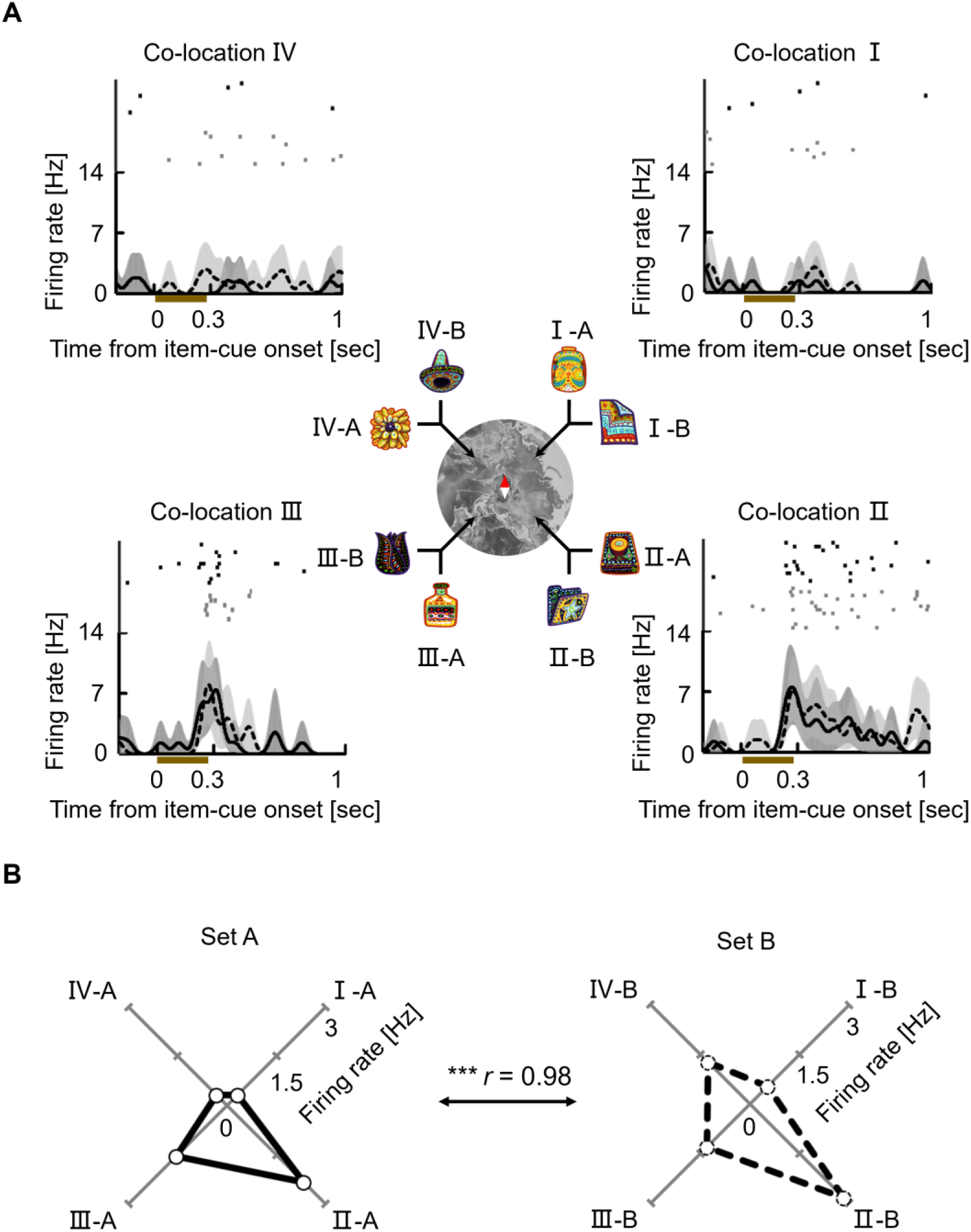
A PRC neuron showing the co-location effect on item-cue selective activities. **(A)** Solid lines and dashed lines indicate spike density functions (SDFs) in trials with item-cues from the stimulus sets A and B, respectively. Dark and light gray shading, 90% confidence interval of 10,000 bootstraps (see Methods) for the stimulus sets A and B, respectively. Black and gray dots, raster plots for the stimulus sets A and B, respectively. Brown bar, presentation of the item-cue. **(B)** Mean discharge rates of the neuron during the item-cue period for each item. *r*, correlation coefficient. Asterisk indicates the result of a two-tailed permutation test: *P* = 0.0024***.

We evaluated the association effect in each area by calculating the co-location index for each one of the item-cue selective neurons. In addition to the HPC, which showed a high co-location index (median, *r* = 0.89, *P* < 0.0001, two-sided Wilcoxon signed-rank test) in our previous study (Yang and Naya, 2020), both the PRC and the PHC showed significantly positive co-location indices (*r* = 0.63, *P* < 0.0001 in PRC; *r* = 0.63, *P* = 0.0009 in the PHC) (Fig. 4A). In contrast with the striking association effect in the MTL, TE did not show a significantly positive co-location index (*r* = 0.16, *P* = 0.0754), which is consistent with previous lesion studies indicating preferential involvement of TE in the perceptual processing of visual objects and preferential involvement of the MTL (PRC) in mnemonic processing (Buffalo et al., 2000).

**Fig. 4.**
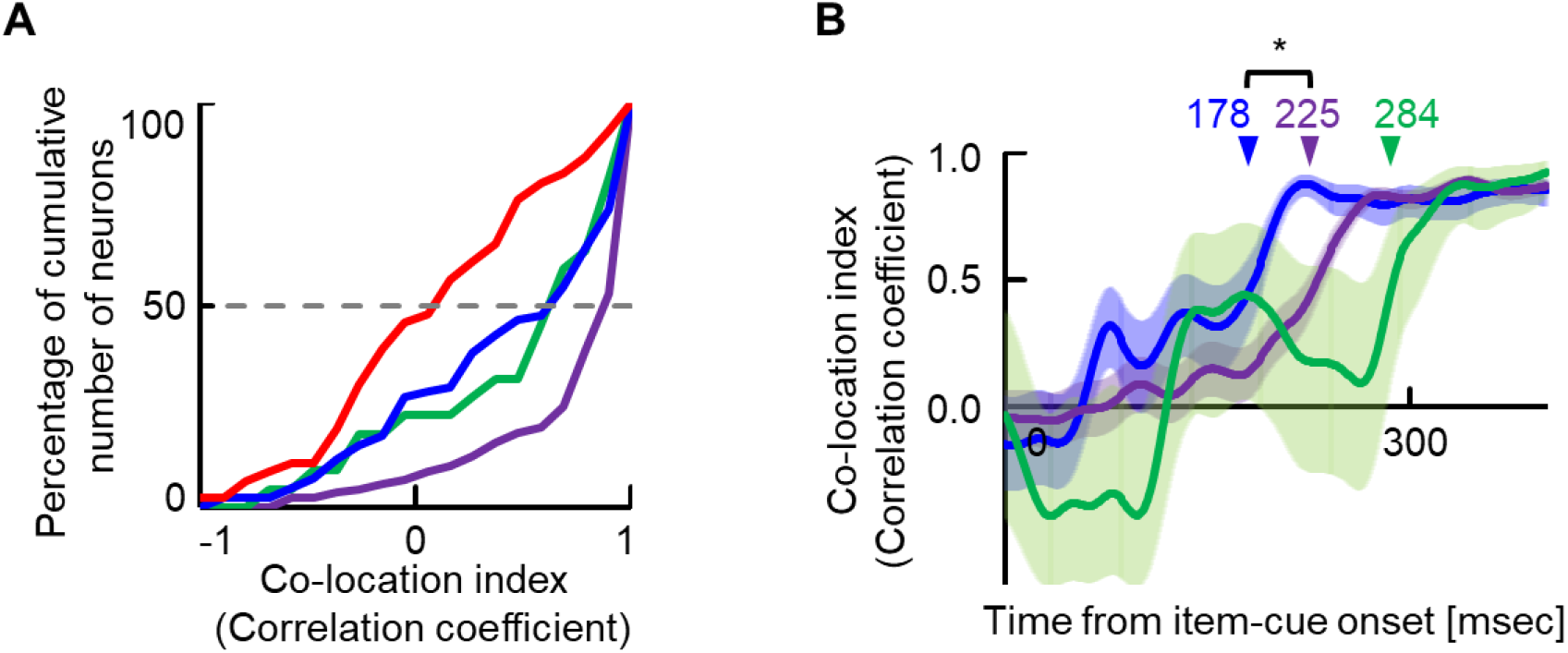
Memory retrieval signal in the MTL. **(A)** Cumulative frequency histograms of co-location indices for item-cue selective neurons in each area. *Red*, TE (n = 47); *blue*, PRC (n = 84); *purple*, HPC (n = 136); *green*, PHC (n = 22). Co-location indices in all MTL areas were significantly positive (*P* < 0.0001, Y = 0, for PRC, *P* < 0.0001, Y = 0, for HPC, *P* = 0.0009, Y = 0, for PHC, two-sided Wilcoxon signed-rank test) and greater than those in TE (*P* = 0.0043, KS = 0.31 for PRC; *P* < 0.0001, KS = 0.59 for HPC; *P* = 0.0056, KS = 0.43 for PHC; Kolmogorov–Smirnov test). **(B)** Time course of co-location index. Lines and shading, mean and standard error of the mean (SEM) across the item-cue selective neurons with high co-location indices (*r* > 0.8) in PRC (*blue*, n = 30), HPC (*purple*, n = 83) and PHC (*green*, n = 8). Arrow, half-peak time. Asterisk indicates the result of a two-tailed permutation test: *P* = 0.0382*.

We next examined which area in the MTL first exhibited the association signal by comparing the time courses of the population-averaged co-location indices among the three MTL areas. To this end, we selected neurons with high co-location indices (*r* > 0.8) and found that the co-location index increased earlier in the PRC than in the HPC and the PHC (Fig. 4B). These results held true for the neurons that were selected at a different threshold (*r* > 0.7, Fig. S4). Considering the dense fiber projections from the TE to the PRC (Saleem and Tanaka, 1996; Suzuki and Naya, 2014), the perceptual information signaling the item-cue in TE may elicit a reinstatement of the item-location associative memory in the PRC, which is consistent with previous physiological studies reporting substantial association effects in the PRC compared with TE under the item-item association paradigm (Koyano et al., 2016; Naya et al., 2001, 2003; Takeda et al., 2015), item-reward association paradigm (Liu and Richmond, 2000; Mogami and Tanaka, 2006), and temporal-order memory paradigm (Eradath et al., 2015; Naya and Suzuki, 2011).

### First-person perspective of the retrieved location

The item-cues elicited item-cue selective activity according to their co-locations, which were determined relative to the map image. However, it is yet to be addressed whether co-location-related activity signaled the retrieved location in relation to a particular spatial context. To answer this question, we compared the responses during the item-cue period with those during the choice-fixation period in which the monkeys fixated on a target location (Fig. 1C). Before the comparison, we examined the representation of task-related information during the choice-fixation period by applying a three-way (item-cue, context-cue, target) ANOVA (*P* < 0.01) for each neuron in the MTL. A substantial number of neurons exhibited differential responses among the target locations (target-selective) in all the MTL areas (6.0% in PRC, 12.1% in HPC, and 10.8% in PHC) (Fig. 5A, Table S1), whereas only a negligible number (∼ 1.5%) of neurons showed item-cue or context-cue selective responses across the three subareas (Fig. 5A, Table S1). These results indicate that the MTL neurons represented the target location where the monkeys presently gazed during the choice-fixation period rather than the information from the preceding task events (i.e., item-cue and context-cue).

**Fig 5.**
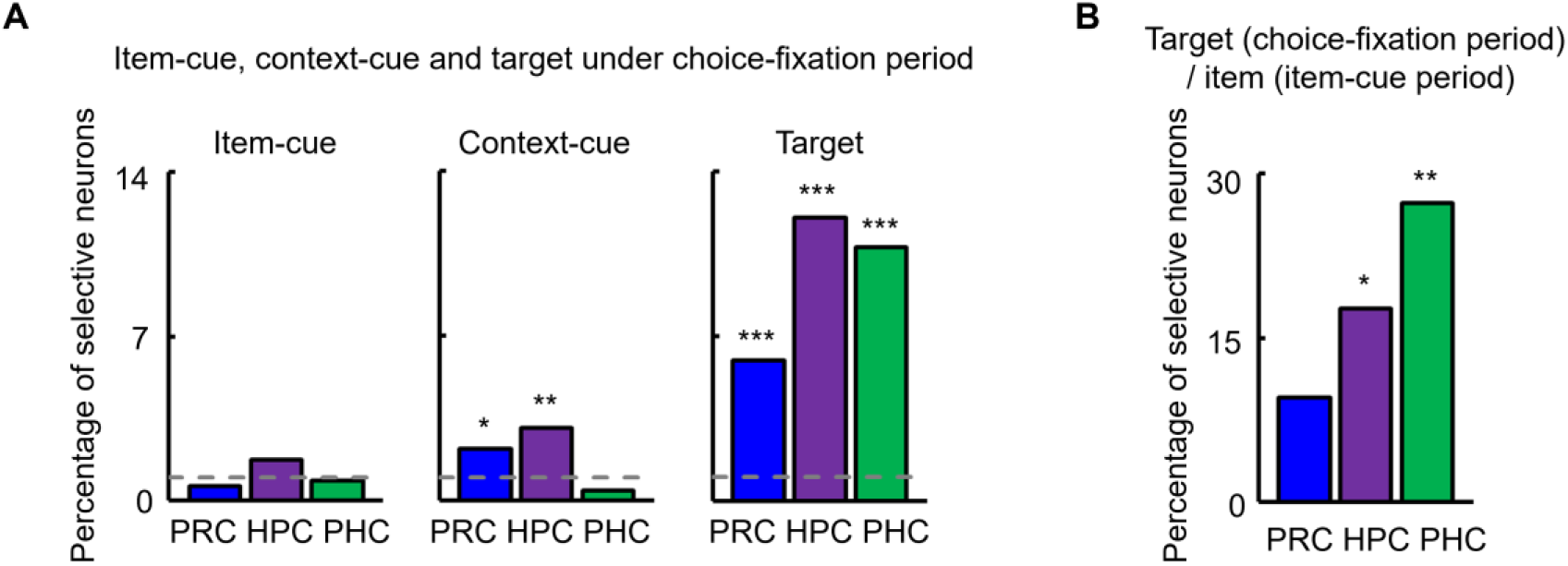
Task-related signals during the item-cue period and choice-fixation period. **(A)** Percentages of item-cue, context-cue, and target-selective neurons out of the recorded neurons during the choice-fixation period. PRC, n = 319; HPC, n = 456; PHC, n = 232. Dashed line, 1.0% chance level. Asterisks indicate results of one-tailed binomial test (probability of null hypothesis = 1.0%): *P* = 0.0432* for context-cue selective neurons in PRC; *P* = 0.0003** for context-cue selective neurons in the HPC; *P* < 0.0001*** for target-selective neurons in each area. **(B)** Percentage of target-selective neurons during choice-fixation period out of item-cue selective neurons (PRC, n = 84; HPC, n = 136; PHC, n = 22). Asterisks indicate results of a χ-square test: *P* = 0.0170*, χ^2^ = 5.7, *d.f* = 1; *P* = 0.0087**, χ^2^ = 6.88, *d.f* = 1.

To compare the responses between the two task periods in each MTL area, we first examined whether neurons exhibiting item-cue selective responses during the item-cue period (i.e., item-cue selective neurons) also showed target-selective responses during the choice-fixation period. We found that the item-cue selective neurons had a significant tendency to show target-selective responses in the HPC (17.6%, *P* = 0.0170, χ^2^ = 5.7, *d.f* = 1, χ-square test) and PHC (27.3%, *P* = 0.0087, χ^2^ = 6.88, *d.f* = 1), but not in the PRC (9.5%, *P* = 0.1075, χ^2^ = 2.59, *d.f* = 1) (Fig. 5B). We confirmed this area difference by considering the strengths (R^2^ values) of item and target selectivity for all recorded neurons (Fig. S5). These results suggest that the neural representations during the item-cue period may be explained by activities for target locations during the choice-fixation period in the HPC and the PHC but not in the PRC.

Next, we compared the preferred locations of neurons during the item-cue period to those during the choice-fixation period. Figure 6 displays the responses of one HPC neuron that showed selective activity during both task periods. This neuron signaled co-location III during the item-cue period, whereas it signaled the bottom-left target location during the choice-fixation period (Figs. 6A and 6B). The bottom-left was assigned as a target location in trials with combinations of “co-location II & 90°,” “co-location III & 0°,” and “co-location IV & -90°” (Fig. 6C). The response patterns to the target locations during the choice-fixation period and those to the co-locations during the item-cue period were positively correlated when the co-locations were assumed to be positioned relative to the 0° context -cue (*r* = 0.99), but not to the -90° context-cue (*r* = -0.27) nor the 90°context -cue (*r* = -0.27) (Fig. S6). These results suggest that the HPC neuron represented the co-location of the item-cue stimulus relative to a particular context-cue during the item-cue period, although the context-cue was not yet presented.

**Fig 6.**
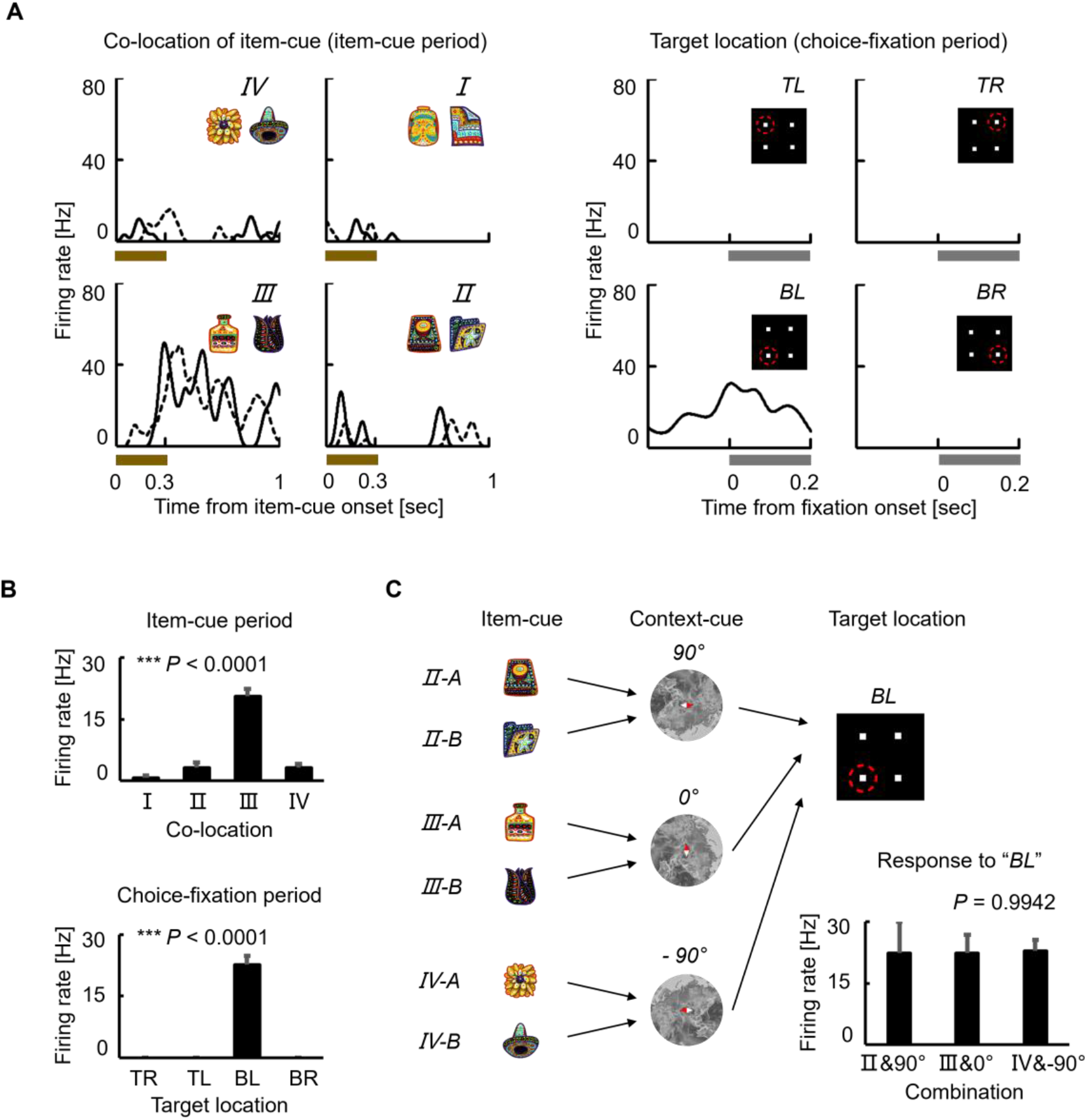
An HPC neuron exhibiting both item selectivity and target selectivity. **(A)** Lines, SDFs for each item-cue during the item-cue period (left) or SDF for each target location during the choice-fixation period (right). I-IV, co-location I-IV. TR, top-right; BR, bottom-right; BL, bottom-left; TL, top-left. **(B)** Mean responses during the item-cue period for each co-location (top) and during the choice-fixation period for each target location (bottom). Error bar, standard error. Significant selectivity for the co-locations (*P* < 0.0001***, *F* (3, 42) = 56.6, one-way ANOVA) and for the target locations (*P* < 0.0001***, *F* (3, 42) = 129.3). **(C)** Combinations of item- and context-cues resulted in the bottom-left (BL) target location. Mean responses during the choice-fixation period for combinations resulted in the BL target location. Error bar, standard error. Responses were not significantly different among the combinations that resulted in the BL target location (*P* = 0.9942, *F* (2, 8) = 0.0058, one-way ANOVA).

We examined this tendency in the population by conducting representational similarity analysis (RSA) using the responses from all the recorded neurons in each area (n = 319, PRC; n = 456, HPC; n = 232, PHC) because only a few neurons showed task-related activity in both task periods in the PRC (n = 8) and the PHC (n = 6). We calculated a correlation coefficient between the *N*-dimensional population vector for the response to each co-location during the item-cue period and that of the response to each target location during the choice-fixation period in each area (Fig. S7). “*N*” was the number of the recorded neurons in each area. According to three orientations (−90°, 0°, or +90°) of the map image, co-locations were combined with particular target locations (e.g., “co-location I” and “top-left” in −90° orientation) to form a total of four combinations in each orientation. The average of the correlations across the four combinations showed a significantly positive value in the HPC (*r* = 0.14 and *P* < 0.0001, two-tailed permutation test) and the PHC (*r* = 0.18, *P* < 0.0001) only when assuming the co-locations on a map image with 0° orientation (Fig. 7). These results suggest that the co-location of the item-cue was represented relative to the 0° map image before the presentation of the context-cue in the HPC and the PHC. The selective conjunction of the co-locations and the 0° map image might be due to the training history in which the monkeys had learned the item-location association under the context-cue with 0° orientation during the initial training (“defult condition”) (Yang and Naya, 2020). In contrast to the HPC and the PHC, a positive correlation was not found for any orientation of the map image in the PRC (*P* = 0.73, 0.77, 0.06 for -90°, 0°, and 90°, respectively). The non-significant result of the RSA in the PRC could neither be explained by the sample size nor the strengths of task-related signal because they were even larger in the PRC than in the PHC (n = 319 in PRC vs. 232 in PHC, sample size; *r* = 0.63 vs. 0.63, co-location index; 26.3% × 6.0% vs. 9.5% × 10.8%, item-cue × target location selective neurons). Together, the comparison of the response patterns between the item-cue and choice-fixation periods suggests that the HPC and the PHC represent the co-locations of the item-cue stimuli in relation to a particular spatial context. Conversely, the PRC might represent co-locations irrespective of their relationship with any particular spatial context.

**Fig 7.**
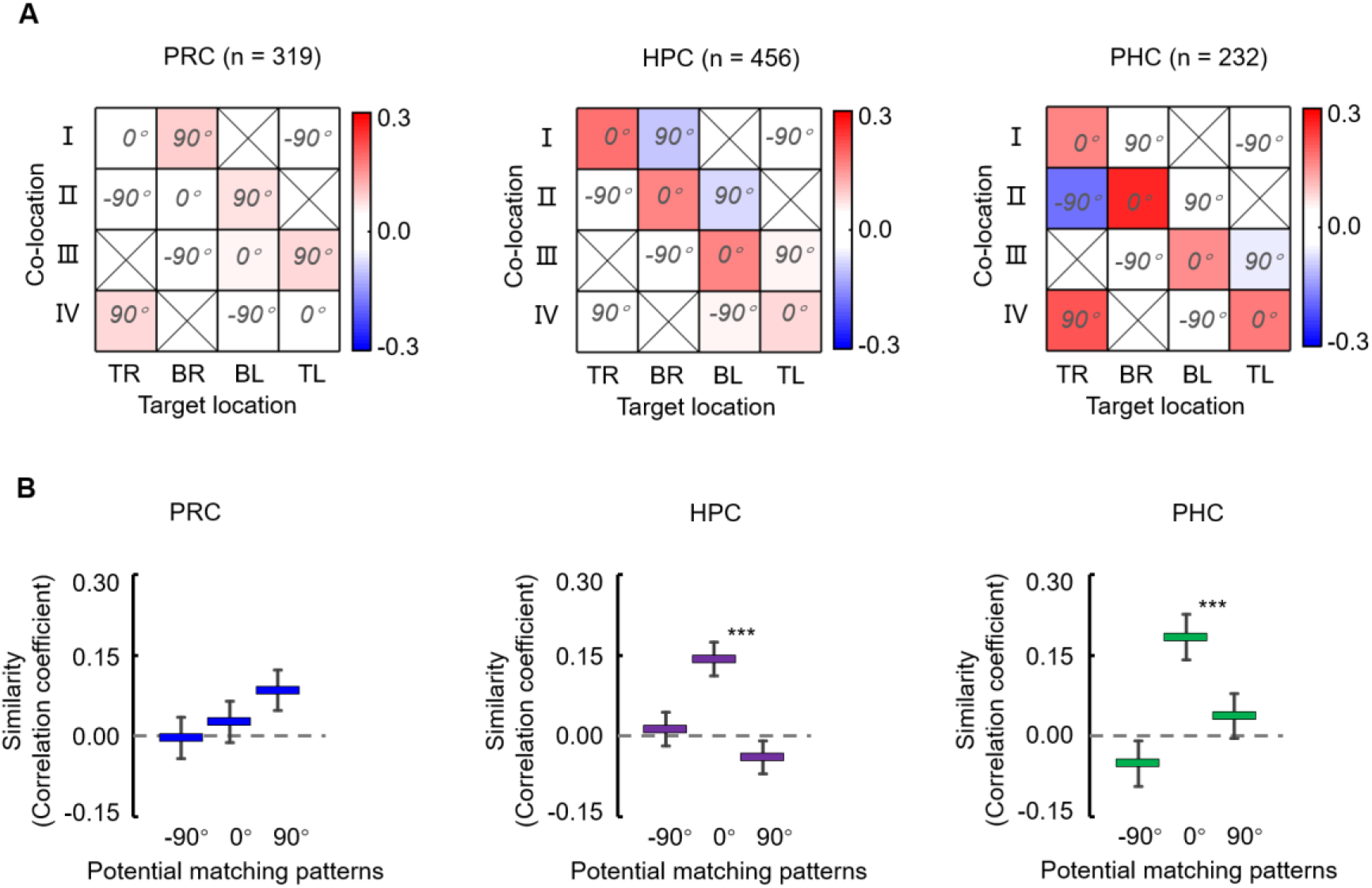
Representational similarity between co-locations and target locations. **(A)** Correlation matrices between population vectors for co-locations of item-cues and those for target locations in each MTL area. The population vectors consist of activity from the recorded neurons in each area (PRC, n = 319; HPC, n = 456; PHC, n = 232). TR, top-right; BR, bottom-right; BL, bottom-left; TL, top-left. **(B)** Representational similarity between co-locations and target locations was estimated as a mean value of correlation coefficients for four combinations of co-locations and target locations corresponding to each of -90°, 0°, and 90° map image conditions. Error bar, standard deviation. Asterisk indicates results of two-tailed permutation test: *P* < 0.0001***.

## DISCUSSION

Using the same ILA task, we previously presented the retrieved co-location information represented in the HPC (Yang and Naya, 2020). However, it remained unclear whether the co-location information was retrieved in the HPC or whether the HPC received a retrieval signal from other upstream brain areas. To address this problem, the present study examined the parahippocampal cortical areas, including the PRC and the PHC, which provide signal input to the HPC. In addition to the HPC, item-cue selective neurons in the PRC and the PHC of the MTL showed significantly correlated responses to pairs of the co-locating items that the monkeys had learned to associate with the same co-locations in advance (Fig. 4A). Among the MTL areas, correlated responses first appeared in the PRC (Fig. 4B). In contrast to the MTL areas, correlated responses were not found in TE, which reportedly provides the PRC with perceptual information of visual objects (Naya et al., 2001; Suzuki and Naya, 2014). These results suggest that PRC was first involved in the memory retrieval process before the HPC. We also examined the representation property of the co-location information in each MTL area and found that the retrieved co-location information was represented in relation to a particular spatial context (i.e., 0° of the map image) in the HPC and the PHC but not in the PRC (Fig. 7). These results suggest that remembering semantic memory may involve neural processing in both the PRC and the HPC (Fig. 8).

**Fig 8.**
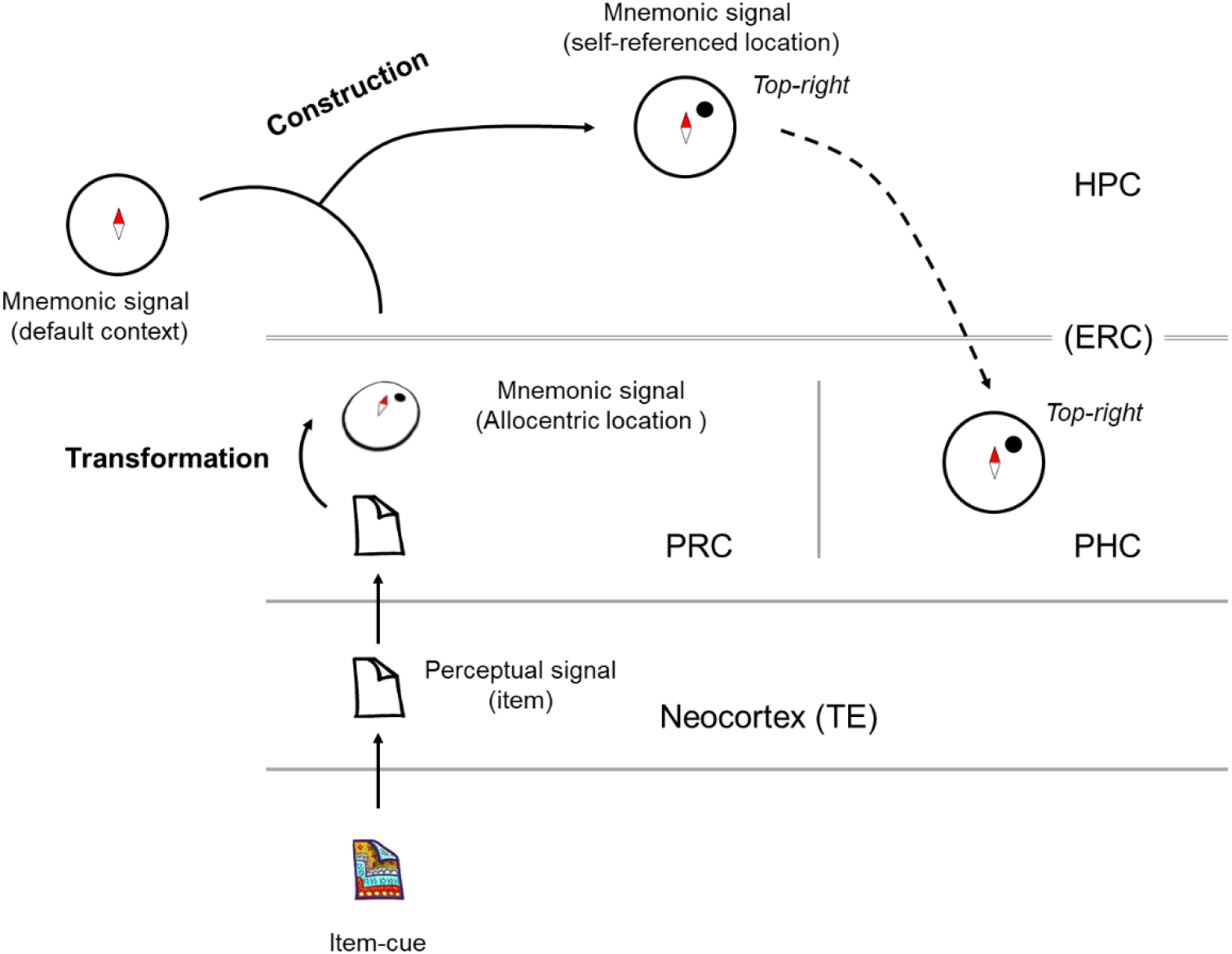
Two-stage recall model of semantic memory. Schematic diagram of neuronal signals in the MTL during the item-cue period in the ILA task. Perceptual signal of the item-cue transmits from the neocortex to the PRC, in which the item information would be transformed to the mnemonically-linked location information, which is represented in an allocentric manner. The retrieval signal would transmit (via the ERC) to the HPC, in which the allocentric location would be combined with the default context to construct the self-reference representation of the retrieved location. The self-reference location signal would spread from the HPC (via ERC) to PHC.

In this study, the effect of item-location associative memory was assessed by correlated responses to pairs of co-locating items. Similar to the present results, an increase in correlated responses to pairs of items from TE to the PRC was also found in previous studies using the pair-association paradigm (Miyashita, 2019; Naya, 2016a; Naya et al., 2003), which examined a direct association between items (Murray et al., 1993; Sakai and Miyashita, 1991). This study using the ILA task indicated that the PRC also served as an indirect linkage between items via their co-locations on the map image. Secondly, to examine the effect of item-location associative memory, we compared activities during the item-cue period with those during the choice-fixation period. This study showed that neurons in the MTL areas exhibited selective responses to target locations where the monkeys positioned their gaze. These target-selective responses during the choice-fixation period could be explained by locations relative to “multiple spatial reference frames” (Meister and Buffalo, 2018), including head-center, external landmarks (e.g., computer screen) as well as to-be-retained context-cue in mind which could be represented in either head-center or external landmark frame. This ambiguity makes it difficult to distinguish the target-selective activity in the self-referenced frame from that in the allocentric frame. However, the gaze-position-dependent activity during the choice-fixation period still seems to reflect the first-person perspective because it depended on what the subjects perceived at that time.

The item-location associative memory isolated from a particular spatial context (e.g., the map image with 0° orientation) in the PRC is consistent with preceding literature suggesting its involvement in semantic memory (Davies et al., 2004; Davies et al., 2009; Lee et al., 2006; Naya et al., 2001; Suzuki and Naya, 2014; Wright et al., 2015) as well as the allocentric representations of semantic memory (Buzsáki et al., 2022; Buzsaki and Moser, 2013). The isolation from the particular context may also explain the previous studies suggesting a selective involvement of the PRC in familiarity rather than in recollection (Aggleton et al., 2012; Eichenbaum et al., 2007). The reactivation of item-location associative memory in the PRC may induce familiarity (Tamura et al., 2017), but the reactivated mnemonic information in the PRC may not provide a first-person perspective that could be re-experienced in mind with a particular spatial context (Buzsáki et al., 2022; Naya, 2016b; Tulving, 2002). Therefore, the location information in the PRC may not be directly coupled with mental imagery as recollection. The poor access of the reactivated item-location associative memory to mental imagery may also explain previous studies suggesting the exclusive processing of object-related information in the PRC (Davachi, 2006; Mayes et al., 2007), although recent physiological studies, including ours, reported space-related information represented by PRC neurons (Ahn and Lee, 2017; Chen and Naya, 2020a, b; Ohnuki et al., 2020; Suzuki and Naya, 2014). Here, we hypothesize that the PRC might be involved in retrieving semantic memory even when it includes the space-related information, although its reinstated spatial contents may not be directly experienced in the mind because of the allocentric representation of semantic memory (Buzsáki et al., 2022).

In contrast to the PRC, the retrieved location was represented in relation to a particular spatial context (0° orientation of the map image) in the HPC and the PHC. An accompaniment of a particular spatial context is consistent with previous studies suggesting the involvement of the HPC in semantic recall with a vivid mental image (Duff et al., 2019; Keane et al., 2020; Lynch et al., 2020), which presumably involves concrete representations from the first-person perspective. Of the two MTL areas, the HPC showed a much larger retrieval signal than the PHC, which was estimated by the proportions of the item-cue selective neurons (29.8% in the HPC vs. 9.5% in the PHC) and their co-location indices (median *r*-value, 0.89 vs. 0.63) (Figs. 2B and 4A). Moreover, the time courses of the co-location index elevated earlier in the HPC than in the PHC (half-peak time, 225 ms vs. 284 ms) (Fig. 4B), which may imply a transmission of the retrieved information from the HPC to the PHC. Together, it may be reasonable to deduce a critical involvement of the HPC rather than the PHC in accompanying semantic recall with a spatial context to represent the memory component in the first-person perspective.

One remaining question is how the retrieved location was accompanied by the particular spatial context during the absence of the map image. Using the same ILA task, we previously showed a constructive process in which the HPC neurons fitted the retrieved location to the spatial context given by the context-cue (Yang and Naya, 2020). Similarly, we hypothesize that before the context-cue presentation, the HPC neurons combined the item-location associative memory and the default context to represent the item-location in the first-person perspective (Fig. 8). Considering the anatomical hierarchy of the PRC linking the ventral pathway and the HPC for object-related information (Chen and Naya, 2021; Kravitz et al., 2013; Suzuki and Naya, 2014), and the earlier rise of the co-location index in the PRC than in the HPC after the item-cue presentation, it seems reasonable to interpret that the HPC neurons received the allocentric item-location from the PRC via the ERC and fit it to the default context in the absence of the map image (Fig. 8). Taken together, we propose a two-stage recall process of semantic memory. In this model, the PRC and the HPC would contribute to the recall of semantic memory in distinct but complementary manners; the PRC might serve for semantic recall in the allocentric frame, while the HPC might serve for constructing the first-person perspective.

## ACKNOWLEDGMENTS

We thank S. Xue for expert animal care. We thank J. Gao, W. Men, G. Yang, and the National Center for Protein Sciences at Peking University for assistance with MRI scanning. We thank D. Lanham for providing the source images of the main stimulus set.

## AUTHOR CONTRIBUTIONS

Y.N. designed the experiments. C.Y. performed the experiments. C.Y. and Y.N. analyzed data and wrote the manuscript.

## METHODS

### KEY RESOURCES TABLE

| REAGENT or RESOURCE | SOURCE | IDENTIFIER |
| --- | --- | --- |
| Experimental Models: Organisms/Strains |  |  |
| Rhesus macaque (*Macaca mulatta*) | Beijing Institute of Xieerxin Biology Resource | http://www.xexbio.com/ |
| Software and Algorithms |  |  |
| MATLAB | MathWorks | http://www.mathworks.com |
| Plexon offline sorter | Plexon | http://plexon.com |
| Other |  |  |
| Microelectrode | AlphaOmega | https://www.alphaomega-eng.com |
| Microprobe | NeuroNexus | https://neuronexus.com |

### RESOURCE AVAILABILITY

#### Lead contact

Further information and requests for reagents and resources should be directed to and will be fulfilled by the lead contact, Yuji Naya.

#### Materials availability

This study did not generate new unique reagents.

#### Data and code availability

The data and code that support the findings of the current study are available from the lead contact on reasonable request.

### EXPERIMENTAL MODEL AND SUBJECT DETAILS

#### Animals

Two adult male rhesus monkeys (*Macaca mulatta*; 6.0–9.0 kg) were included in this study. All procedures and treatments were performed in accordance with the National Institutes of Health (NIH) Guide for the Care and Use of Laboratory Animals and approved by the Institutional Animal Care and Use Committee (IACUC) of Peking University (Psych-YujiNaya-1).

### METHOD DETAILS

#### Behavioral task

Two monkeys were trained on an ILA task (Fig. 1) and performed both the training and recording sessions under dim lights. The task was initiated by the monkey fixating on a white square (0.4° visual angle) in the center of a display for 0.5 s. Eye position was monitored by an infrared digital camera with a 120-Hz sampling frequency (ETL-200, ISCAN). Subsequently, an item-cue (diameter, 3.4°) and context-cue (diameter, 28.5°) were sequentially presented for 0.3 s each with a 0.7 -s interval. After a 0.7-s delay interval, four white squares (0.4°) were presented as the choice stimuli an equidistance from the center (6°). One of the squares was a target, whereas the other three were distracters. The target was determined using a combination of the item-cue and the context-cue stimuli. The monkeys were required to saccade one of the four squares within 0.5 s. If they made the correct choice, four to eight drops of water were given as a reward. The trial was terminated without a reward when the monkeys failed to maintain their fixation (typically less than 2° from the center) before the presentation of the choice stimuli. The monkeys were trained to associate two sets of four visual stimuli (item-cues) with four particular locations relative to the context-cue image that was presented at a tilt (with an orientation from -90° to 90°) before the recording session commenced. To prevent the monkeys from learning to associate each combination of the item-cue and context-cue with a particular target location, the orientation of the map image was randomized at a step of 0.1°, which increased the number of combinations (8 × 1,800) and would make it difficult for the monkeys to learn all the associations among the item-cues, context-cues, and target locations directly. During the recording session, the item-cue was pseudorandomly chosen from the eight well-learned visual items, and the orientation of the context-cue was pseudorandomly chosen from among five orientations (−90°, - 45°, 0°, 45°, and 90°) in each trial, resulting in 40 (8 × 5) different configuration patterns. We trained both monkeys using the same stimuli but with different item-location association patterns. All the stimulus images were created using Photoshop (Adobe). Both monkeys performed the task correctly (chance level = 25%) at rates of 80.5% ± 8.4% (mean ± standard deviation; Monkey B, n = 435 recording sessions) and 95.1% ± 5.8% (Monkey C, n = 416 recording sessions).

#### Electrophysiological recording

The monkeys were each implanted with a head post and a recording chamber under aseptic conditions, using isoflurane anesthesia, following the initial behavioral training. We used a 16-channel vector array microprobe (V1 X 16-Edge; NeuroNexus) or a single-wire tungsten microelectrode (Alpha Omega) to record single-unit activity, which was advanced into the brain using a hydraulic Microdrive (MO-97A; Narishige) (Naya and Suzuki, 2011). The microelectrode was inserted through a stainless-steel guide tube positioned in a customized grid system in the recording chamber. Neuronal signals for single units were collected (low-pass, 6 kHz; high-pass, 200 Hz) and digitized (40 kHz) (AlphaLab SnR Stimulation and Recording System, Alpha Omega). These signals were sorted using an offline sorter (Plexon). An average of 87 trials were tested for each neuron (n = 1,175). The placement of the microelectrodes into the target areas was guided by individual brain atlases from MRI scans (3T, Siemens). Individual brain atlases were also constructed based on the electrophysiological properties around the tip of the electrode (e.g., gray matter, white matter, sulcus, lateral ventricle, and bottom of the brain). The recording sites were estimated by combining the individual MRI and physiological atlases (Naya et al., 2017).

The recording sites covered 3–19 mm anterior to the interaural line (monkey B, left and right hemisphere; monkey C, right hemisphere; Fig. 2A). The recording sites in the HPC appeared to cover all its subdivisions (i.e., the dentate gyrus, CA3, CA1, and subicular complex) (Yang and Naya, 2020). The recording sites in the PRC appeared to cover areas 36 from the fundus of the rhinal sulcus to the medial lip of the anterior middle temporal sulcus (amts) (Naya et al., 2003; Saleem and Tanaka, 1996; Suzuki and Amaral, 1994). The recording sites in the PHC appeared to cover area TFl (Suzuki and Amaral, 2003). The recording sites in the TE were limited to the ventral area, including both banks of the amts.

### QUANTIFICATION AND STATISTICAL ANALYSIS

All neuronal data were analyzed using MATLAB R2020a (MathWorks) with custom-written programs, including the statistics toolbox. The permutation tests were performed with 10,000 shuffle iterations.

#### Classification of task-related neurons

For the item-cue period, we calculated the mean firing rates of eight consecutive 300-ms time-bins moving in 100-ms steps, from 0 to 1000-ms after item-cue onset, across all correct trials. We evaluated the effects of “item” for each neuron using one-way ANOVA with the eight item-cue stimuli as the main factor (*P* < 0.01, *Bonferroni correction* for eight-analysis time windows). We refer to neurons with significant item effects during any of the eight-analysis time windows as item-cue selective neurons. For the choice-fixation period, we calculated the mean firing rates of the 200-ms choice-fixation period when the subject fixated on the target location across all correct trials with -90°, 0°, and 90° context-cues. We evaluated the effects of “item,” “context,” and “target location” for each neuron by using three-way ANOVA with the eight item-cue stimuli, three context-cue orientations, and four target locations as main factors (*P* < 0.01).

#### Display of example neurons

To show the activity time course for an example of an item-selective neuron, a spike density function (SDF) was calculated using only correct trials and was smoothed using a Gaussian kernel with a sigma of 20 ms (Figs. 3A, S2, and S3). The deviation in the response to each item-cue was evaluated using bootstrap resampling. For each correct trial, we smoothed the response across time using a Gaussian kernel with a sigma of 20 ms. For each item-cue, bootstrap data samples and the time courses of mean responses were generated 10,000 times. A 90% confidence interval of the 10,000 bootstraps was plotted. Only SDFs were shown during the item-cue and choice-fixation periods as an example of item-cue selective neurons exhibiting target-selective responses during the choice-fixation period (Fig. 6A).

#### Item-location associative effect during the item-cue period

We examined the item-location associative effect on item-selective activity by calculating the Pearson correlation coefficients between the responses to co-location items. For each item-cue selective neuron, we calculated mean firing rates for each 300-ms time-bin, moving by 100 ms during the item-cue period for each correct trial (eight bins in total). We subsequently performed a one-way ANOVA and calculated the grand mean for each item stimulus across correct trials for each bin. For the bins that showed a significant (*P* < 0.01, *Bonferroni correction* for eight-analysis time windows) item effect, we calculated the correlation coefficients between the mean firing rates of the co-location items. After which, we averaged *Z*-transformed values of the correlation coefficients across time-bins for each neuron. Finally, the average value was transformed into *r* values (i.e., the co-location index).

To examine the time courses of the retrieval signal in each area, we calculated the correlation coefficients for each 100-ms time-bin moving by 1 ms during the item-cue period for each item-cue selective neuron with high correlation coefficients. We then averaged the correlation coefficients using *Z*-transformation for each time-bin across neurons for each area. The half-peak time was defined as the time from the item-cue onset to the instant when the population-averaged correlation coefficient (*r*-value) in each area, reached 50% of its peak rise from 0.

#### Relationships between retrieved co-locations and target locations

The RSA was performed using data from the -correct trials (with -90°, 0°, and 90° context-cues) of all the recorded neurons. For each neuron, we first averaged the responses during the item-cue period (60–1,000 ms period from the item-cue onset) in each trial and calculated a grand mean across trials for each of the four co-locations. For each area, an *N*-dimensional population vector was prepared for each co-location. “*N*” was the number of the recorded neurons in the area. The *i*-th element of the population vector indicated the average firing rate of the *i*-th neuron for the co-location. Four population vectors were generated for the responses during the item-cue period. We also prepared four population vectors indicating the responses to the four target locations during the choice-fixation period (0 to 200 ms from the onset of fixation on the target location) in each area. Each of the -90°, 0°, and 90°context -cues was assigned four particular combinations between co-locations and target locations (Fig. S7). We calculated the Pearson correlation coefficients between the population vectors for a co-location and a corresponding target location in each combination (e.g., ‘co-location Ⅰ’ and ‘top-left’ for -90°). After the Z-transformation, the correlation coefficients were averaged across the four combinations and reversed to *r* value.

## SUPPLEMENTARY FIGURES

**Fig. S1.**
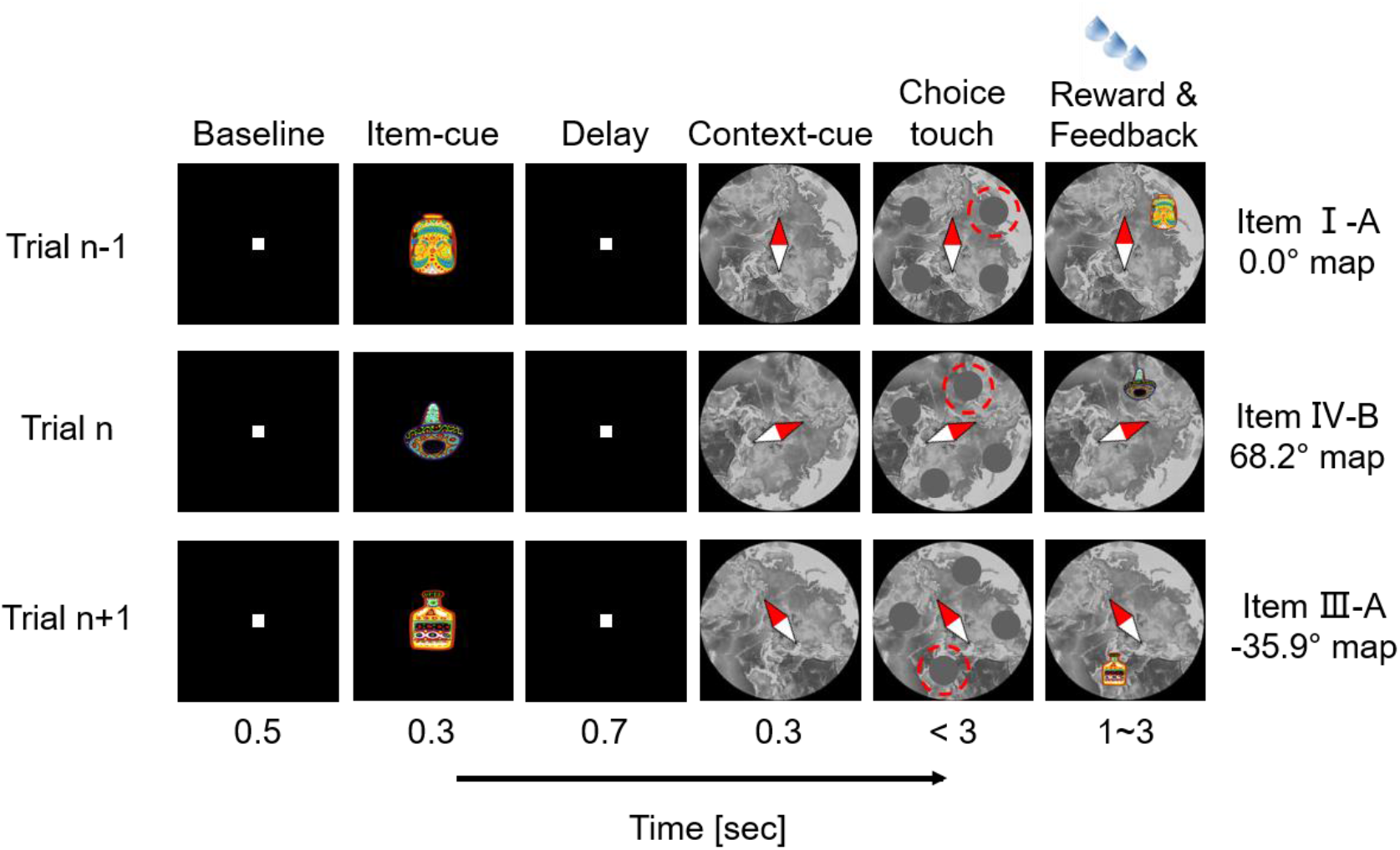
Example trials of the item-location association task during the training phase using a touchscreen. An item-cue and a context-cue were sequentially presented in each trial. The item-cue was chosen randomly from the eight visual items, and the context-cue was presented with a randomly chosen orientation from −90° to 90° in a 0.1° step. Monkeys had to make a choice by touching the target location (red dashed circle) according to the two cues. A successful trial was rewarded with juice paired with feedback showing the associated location of the item-cue on the context-cue. Relative sizes of the stimuli were magnified for display purposes.

**Fig. S2.**
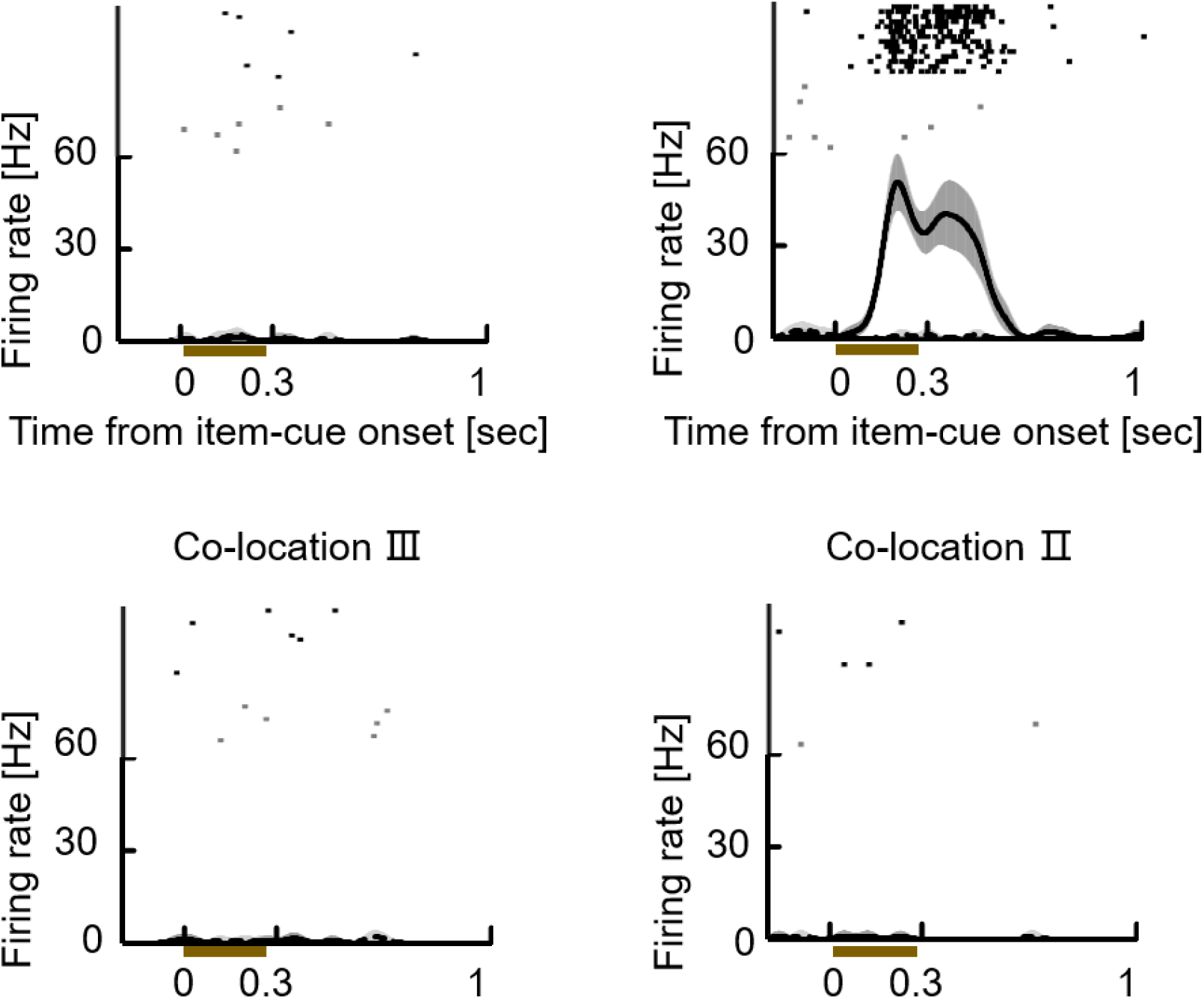
A TE neuron showing the item-cue-selective activities. Following a similar format as Fig. 3A. Solid lines and dashed lines indicate spike density functions (SDFs) in trials with item-cues from the stimulus sets A and B, respectively. Dark and light gray shading, 90% confidence interval of 10,000 bootstraps for the stimulus sets A and B, respectively. Black and gray dots, raster plots for the stimulus sets A and B, respectively. Brown bar, presentation of the item-cue. *P* < 0.0001, *F* (7,115) = 153.41, one-way ANOVA. Co-location index *r* = 0.16, Pearson correlation; *P* = 0.87, two-tailed permutation test.

**Fig. S3.**
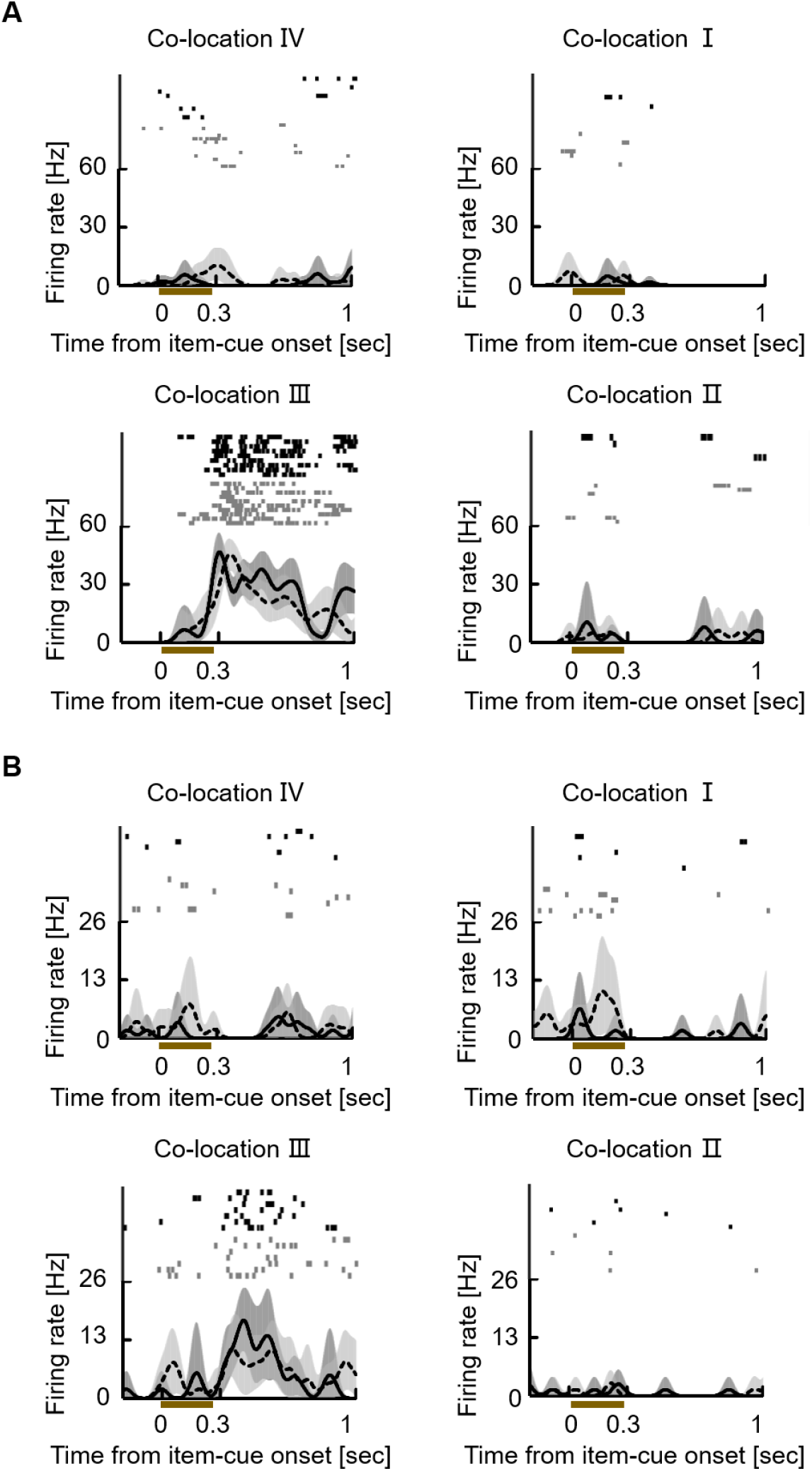
Neurons showing the co-location effect on the item-cue-selective activities. **(A)** An example neuron from HPC. *P* < 0.0001, *F* (7,70) = 45.09, one-way ANOVA. Co-location index *r* = 0.99, Pearson correlation; *P* = 0.0002, two-tailed permutation test. The same format as Fig. S2. **(B)** An example neuron from PHC. *P* < 0.0001, *F* (7,57) = 15.74, one-way ANOVA. Co-location index *r* = 0.99, Pearson correlation; *P* = 0.0004, two-tailed permutation test.

**Fig. S4.**
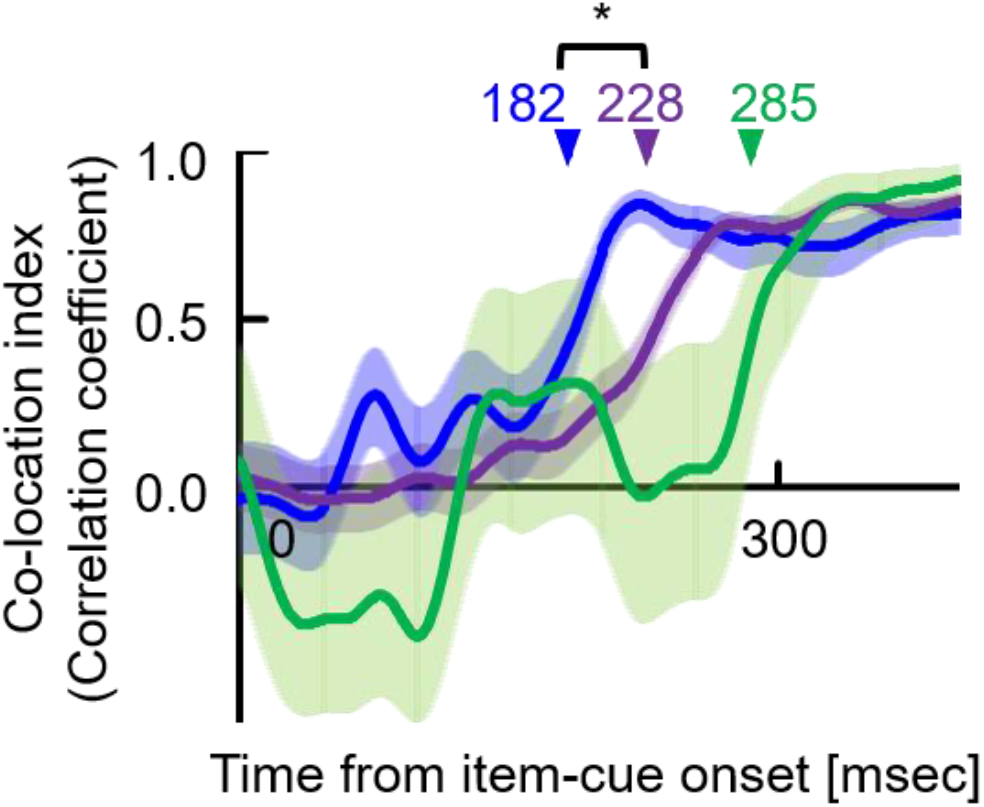
Time course of co-location index using ‘*r* > 0.7’ as a criterion. The time courses of co-location index for the item-cue selective neurons with high co-location indices (*r* > 0.7) in PRC (blue, n = 38), HPC (purple, n = 102) and PHC (green, n = 9). The formats are the same as those in Figure 4B. Asterisk indicates the result of a two-tailed permutation test: *P* = 0.0114*.

**Fig. S5.**
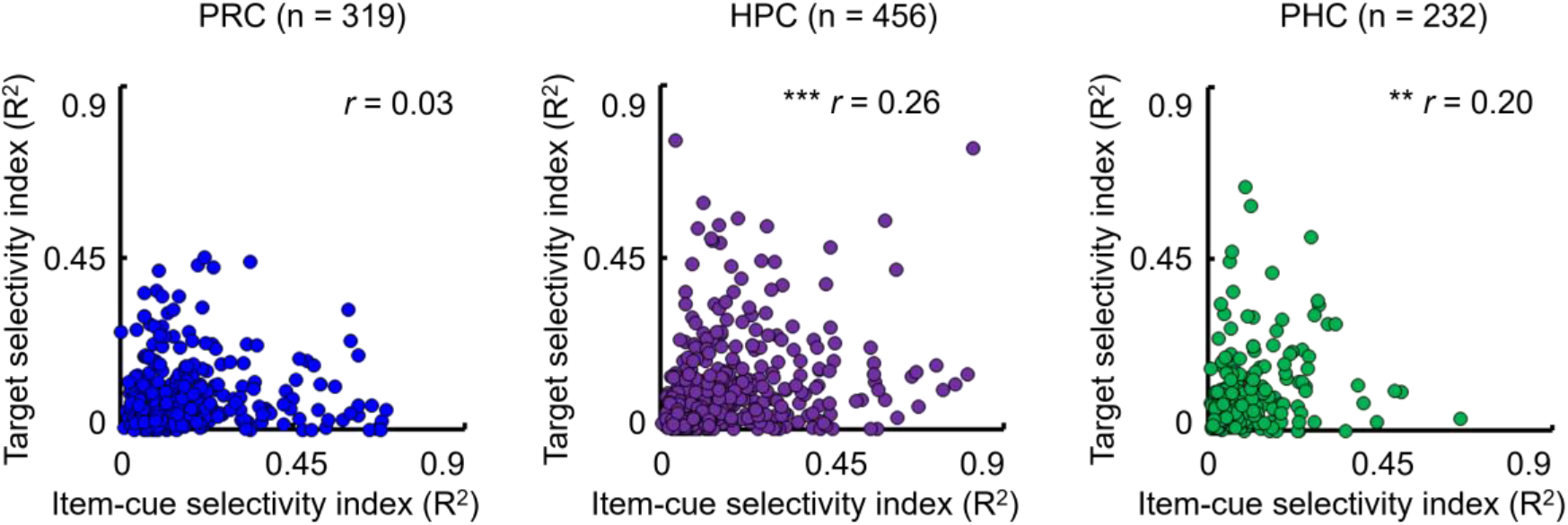
Strengths of item selectivity and target selectivity in the MTL. Item selectivity index, the R^2^ value of item effect from one-way ANOVA test during the item-cue period. Target selectivity index, the R^2^ value of target location effect from three-way ANOVA test during the choice-fixation period. Each dot indicates one neuron. Asterisk indicates the results of a two-tailed permutation test: HPC, *P* < 0.0001***; PHC, *P* = 0.0084**.

**Fig. S6.**
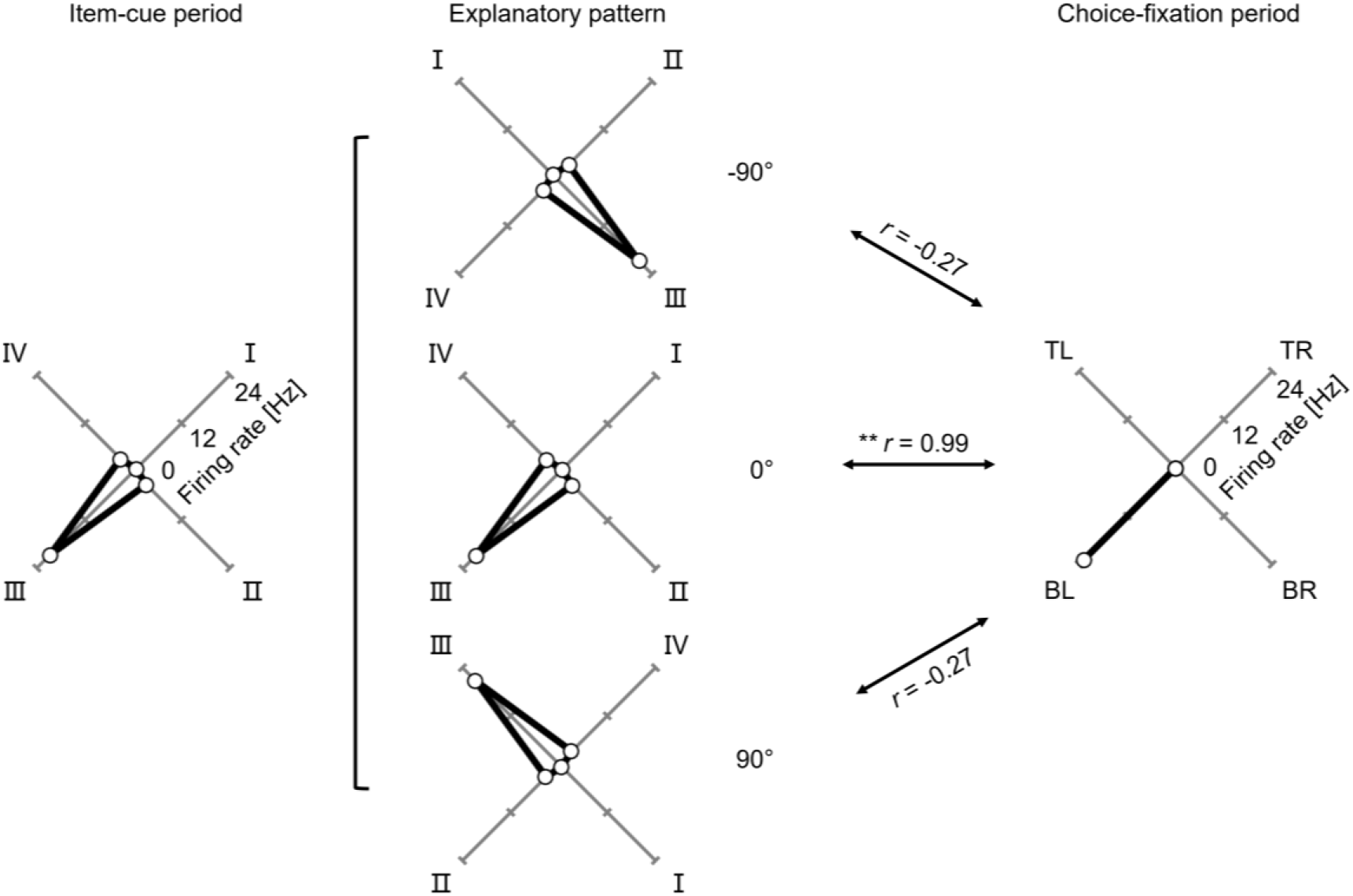
Correspondences between co-locations and target locations under three potential patterns. Explanatory pattern, the co-locations were assumedly positioned relative to a -90°, 0°, or 90° context -cue. *r*, the similarity between response patterns to the co-locations during the item-cue period and those to the target locations during the choice-fixation period. I-IV, co-location I-IV. TR, top-right; BR, bottom-right; BL, bottom-left; TL, top-left. Asterisks indicate the results of a two-tailed permutation test: *P* = 0.0074**.

**Fig. S7.**
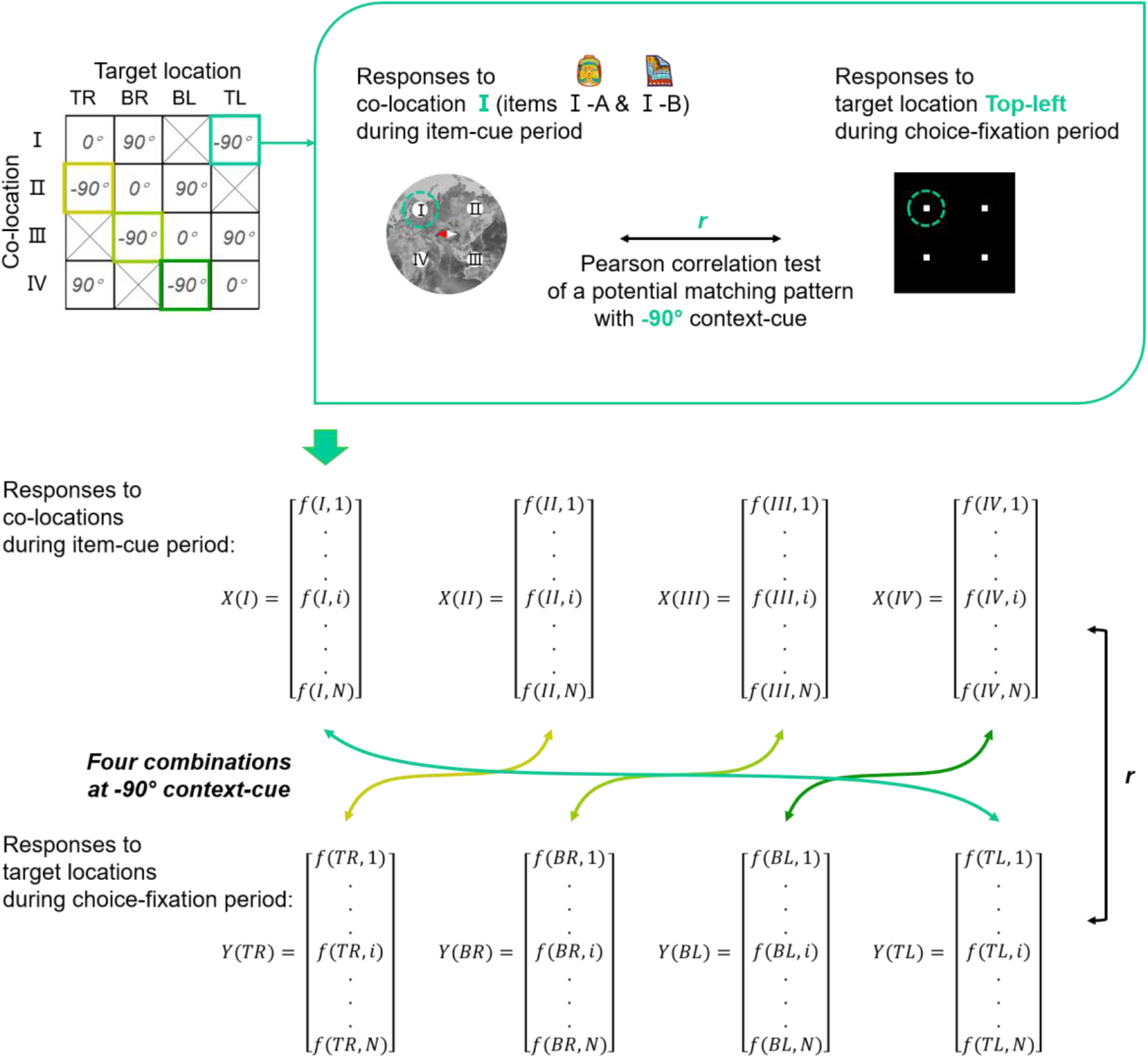
A schematic illustration of RSA. *X(co-location)*, *N*-dimensional population vector for each co-location. *f (co-location, i)*, the average firing rate of the *i*-th neuron for the co-location during the item-cue period. *N*, number of recorded neurons in the area. *Y(target)*, population vector for each target location. *f (co-location, i)*, the average firing rate of the *i*-th neuron for the target location during the choice-fixation period. *r*, Pearson correlation coefficient between *X(co-location)* and *Y(target)*. *-90°*, *0°*, or *90°*, explanatory pattern, the co-locations were assumedly positioned relative to a -90°, 0°, or 90° context-cue.

**Table S1.** Numbers of task-related neurons. Numbers of neurons showing item effect during the item-cue period (*P* < 0.01, one-way ANOVA) and those showing the item, context, and target effects during the choice-fixation period (*P* < 0.01, three-way ANOVA). “Item” indicates an item effect during the item-cue period or choice-fixation period. “Context” indicates a context effect during the choice-fixation period. “Target” indicates a target location effect during the choice-fixation period.

|  |  | *Item-cue period* | *Choice-fixation period* |  |  |
| --- | --- | --- | --- | --- | --- |
|  | **Recorded** | **Item** | **Item** | **Context** | **Target** |
| *Total* |  |  |  |  |  |
| **TE** | 168 | 47 | 2 | 3 | 12 |
| **PRC** | 319 | 84 | 2 | 7 | 19 |
| **PHC** | 232 | 22 | 2 | 1 | 25 |
| **HPC** | 456 | 136 | 8 | 14 | 55 |
| *Monkey B* |  |  |  |  |  |
| **TE** | 98 | 36 | 2 | 1 | 8 |
| **PRC** | 168 | 29 | 1 | 2 | 14 |
| **PHC** | 70 | 7 | 1 | 0 | 7 |
| **HPC** | 247 | 66 | 5 | 7 | 38 |
| *Monkey C* |  |  |  |  |  |
| **TE** | 70 | 11 | 0 | 2 | 4 |
| **PRC** | 151 | 55 | 1 | 5 | 5 |
| **PHC** | 162 | 15 | 1 | 1 | 18 |
| **HPC** | 209 | 70 | 3 | 7 | 17 |

